# Morphogenesis of giant telomeric nuclear bodies in cancer cells with alternative lengthening of telomeres

**DOI:** 10.1101/604959

**Authors:** Yuu Arimasu, Masachika Fujiwara, Jun Ishii, Tomohiro Chiba, Yuichi Terado, Yukiko Shishido-Hara, Hiroshi Kamma

## Abstract

Some cancer cells lengthen their telomeres by alternative lengthening of telomeres (ALT); these are referred to as ALT cancer cells and do not express telomerase. The ALT mechanism involves the elongation of telomeric DNA repeats by homologous recombination. In interphase nuclei of ALT cancer cells, giant telomeres can be specifically observed by fluorescence *in situ* hybridization (FISH) to detect telomeric repeats, and they are co-localized with promyelocytic-leukemia nuclear-bodies (PML-NBs). However, it how large telomeres specific to ALT and how they form a structural relation with PML-NBs. We refer to giant telomeres specific to interphase ALT cancer cells giant-telomeric nuclear-bodies (GT-NBs). To quantitatively define GT-NBs, we performed telomeric FISH of both interphase nuclei and metaphase chromosomes and analyzed telomere sizes by integrated FISH signals. The distributions of telomere sizes in telomerase-positive cells were similar in interphase nuclei and chromosomes. However, the distribution of telomere sizes in ALT cancer cells differed between interphase nuclei and chromosomes. Giant telomeres that were larger than those at chromosomal ends were only observed in interphase nuclei of ALT cancer cells. Accordingly, GT-NBs could be quantitatively defined as larger than the maximum size of telomeres at chromosomal ends. Furthermore, ALT cancer cells demonstrated fewer telomeric signals in interphase nuclei than in chromosomes. These findings indicate that GT-NBs could be formed by the aggregation of two or more telomeres at chromosomal ends. Furthermore, super-resolution microscopy showed that GT-NBs contain one or two PML-NBs. GT-NBs are considered aggregates of telomeres and could contain multiple sites of homologous recombination accompanied by PML-NBs. These findings may contribute to the development of therapeutic approaches for ALT cancer. (251 words)

## Introduction

A telomere is a DNA–protein complex situated at each chromosomal end composed of telomeric DNA with up to 10 kb of TTAGGG repeats and DNA-binding proteins [1-3]. Most cancer cells express telomerase to lengthen telomeric DNA, and these are referred to as telomerase-positive cancer cells (Tel (+) cancer cells) [4-6]. Nevertheless, 10%– 15% of human cancers exhibit little telomerase activity; in these cells, telomeric DNA is elongated by an alternative mechanism [6-9], termed alternative lengthening of telomeres (ALT), involving homologous recombination between telomeric DNAs [5-11]. Cancer cells that lengthen telomeres by the ALT mechanism are referred to as ALT cancer cells. Telomerase inhibitors are not expected to be effective anticancer therapies in ALT cancer. Telomeres of ALT cancer cells vary from 3 to 20 kb but are generally longer than those of Tel (+) cancer cells [3-4]. Telomeres can be detected by fluorescence *in situ* hybridization (FISH) and quantified by integrated brightness fluorescence intensities for telomere signals, reflecting the amount of telomeric DNA. The signals of giant telomeres are observed by FISH in interphase nuclei of ALT cancer cells [4,12-15]. However, it is not clear how large telomeres are specific to ALT cancer cells and how they form in their interphase nuclei. In the present study, we refer to ALT-specific giant telomeres as “giant-telomeric nuclear-bodies (GT-NBs)” and attempted to define GT-NBs in a quantitative manner and to clarify their morphogenesis.

Promyelocytic leukemia nuclear bodies (PML-NBs) are intranuclear structures containing the PML protein; they were first discovered in patients with acute promyelocytic leukemia [13,16-19]. Typically measuring around 0.2–1.0 nm, PML-NBs assume globular structures with doughnut-shaped cross-sections. PML-NBs are involved in various cellular functions, such as the regulation of the cell cycle, aging, and apoptosis [13,16-19]. PML-NBs adhere to chromosomes by binding to proteins, such as death domain-associated proteins, later in the S phase and play a role in the reconstruction of chromatin by binding to chromosomal proteins, as observed in X-linked α-thalassemia/mental retardation (ATRX) syndrome in the G2 phase. Finally, PML-NBs detach from chromosomes during metaphase [20].

Some telomeres are co-localized with PML-NBs in ALT cancer cells, but this does not hold true for Tel (+) cancer cells [13,21]. Telomeres co-localized with PML-NBs are referred to as ALT-associated PML-NBs (APBs) because they are involved in the ALT mechanism [10,12-13,21-22]. So, are APBs are the same nuclear structure as ALT-specific GT-NBs? In order to clarify their relationship, we investigated the ultra-structural relation between GT-NBs and PML-NBs using a super-resolution microscopy whose resolution is 20 times higher than a fluorescence microscopy [23-24].

## Materials and Methods

### Cultured cells

HeLa and TTA-2 cells were used as Tel (+) cancer cells, and H295R, U2OS, and SK-LU1 cells were used as ALT cancer cells. All cultured cell lines were purchased from the American Type Culture Collection (Manassas, VA, USA). U2OS cells are widely used in ALT research, but their chromosomes are unstable. H295R cells have a stable number of chromosomes, making them suitable for morphological analyses.

For HeLa, TTA-2, U2OS, and SK-LU1 cells, Dulbecco’s modified Eagle’s medium (DMEM) (Sigma-Aldrich, St. Louis, MO, USA) was used as the base medium, to which 10% fetal calf serum (FCS) (GIBCO, Gaithersburg, MD, USA), 100 IU/mL penicillin, and 100 µg/mL streptomycin (Sigma-Aldrich) were added. For H295R cells, DMEM and F12 Ham’s medium (Sigma-Aldrich) were mixed in a 1:1 ratio to prepare a base medium, to which 2.5% FCS, 100 IU/mL penicillin, 100 µg/mL streptomycin, selenious acid, and retinoic acid were added [4]. Cultured cells were incubated in a carbon dioxide incubator at 37°C with 5% CO_2_.

### Preparation of chromosome specimens

To analyze telomeric signals during metaphase, chromosomes were prepared. After cultured cells were incubated for 18 h with 10 µg/mL colcemid (GIBCO), samples were fixed with Carnoy’s solution (acetic acid: methanol = 1:3). Subsequently, the sample attached to a slide glass was immediately immersed in a thermostatic bath set to 72°C, thereby evaporating Carnoy’s solution to obtain the specimen.

### Preparation of interphase nuclei

Interphase nuclei of the cultured cells were prepared. Cultured cells were first detached using Trypsin and then collected as a pellet by centrifugation. Subsequently, the sample was fixed by inversion mixing with 4% paraformaldehyde (Wako Pure Chemical Industries, Co. Ltd., Osaka, Japan) at room temperature (25°C) for 30 min before it was washed with phosphate-buffered saline. The sample was then pressure-bonded onto a poly-L-lysine (Sigma-Aldrich)-coated slide glass or cover glass. Because the nucleus is spherical in an interphase cell, the thickness of the specimen makes it difficult to focus on some parts of the nucleus using a microscopic lens. Therefore, high-speed centrifugation was performed to flatten the nucleus for observations on a single plane.

### Telomeric FISH

For each cell line, metaphase chromosomes and interphase nuclei were subjected to 10 µg/mL RNase A treatment at 37°C for 1 h. Subsequently, samples were treated with 1 µg/mL pepsin (Sigma-Aldrich) in 0.5 N hydrochloric acid at 37°C for 10 min before being dehydrated with a dilution series of ethanol. After complete desiccation, samples were subjected to heat treatment at 80°C for 3 min with a telomere-specific peptide nucleic acid probe with Alexa488-(CCCTAA)_3_ (PANAGENE^®^, Daejeon, Korea) and then hybridized at 37°C overnight. After hybridization, samples were washed three times with 2× Saline Sodium Citrate (SSC) Buffer, counterstained with either 4′,6-diamidino-2-phenylindole (DAPI) or propidium iodide (PI), and mounted with 100 mM 2-aminoethanethiol (Wako Pure Chemical Industries). To quantitatively analyze telomeric fluorescence signals, samples were placed on a poly-L-lysine (Sigma-Aldrich)-treated cover glass and were centrifuged at approximately 120,000 × *g*.

### Double fluorescent staining for PML-NBs and telomeres

Cultured cells were immersed in 0.5% Triton X-100 (Sigma-Aldrich) at 4°C for 15 min and then subjected to blocking with 3% albumin (Sigma-Aldrich) at room temperature for 1 h. For primary antibodies (PML-NB; MBL, Nagoya, Japan, 5000-fold dilution) [25] and secondary antibodies (anti-mouse antibodies; Nichirei Histofine SAB-PO(R) Kit), biotin-labeled Alexa 568 was employed as the fluorophore. FISH procedures were followed by immunofluorescent staining. For counterstaining, either DAPI or PI was used.

### Microscopy and image analysis

Fluorescent images were captured using a fluorescence microscope (BX60) (Olympus, Tokyo, Japan) with a 100× objective lens and a digital camera (CoolSNAP ES2 Photometrics) (Roper Scientific, USA). A super-resolution microscope with grand state depletion (SR-GSD M16000B) (Leica, Wetzlar, Germany) was also used [24]. To estimate the size of telomeres and the amount of telomeric DNA, telomeric FISH signals in metaphase chromosomes and interphase nuclei were quantified using Image-Pro^®^(Roper Scientific). Because the size and brightness of a telomeric signal are correlated with the amount of telomeric DNA, the brightness and area of pixels constituting each telomeric signal in the FISH image data were integrated, i.e., the mean brightness and area of all pixels constituting each telomeric signal were multiplied to calculate the integrated value (IV).

## Results

### Telomeric FISH of chromosomes and interphase nuclei

Based on fluorescence microscopy, the telomeric FISH signals for chromosomes of Tel (+) cancer cells (HeLa and TTA-2) and ALT cancer cells (H295R, U2OS, and SK-Lu1) differed from those of their interphase nuclei (Fig 1). The sizes of telomeric signals of chromosomes of all examined cells were small and rather uniform. The sizes of telomeric signals in interphase nuclei of HeLa and TTA-2 cells were also small and similarly uniform. However, those of interphase nuclei of H295R, U2OS, and SK-Lu1 cells were variable, including disproportionately large signals identified as GT-NBs and characteristic of ALT cancer cells. APB and GT-NBs were co-localized with telomeres and PML-NBs. The difference between these two structures is defined by the telomere size. APBs exhibit a range of sizes, including small sizes. GT-NBs are strictly defined by the size of telomeres.

**Fig 1.**
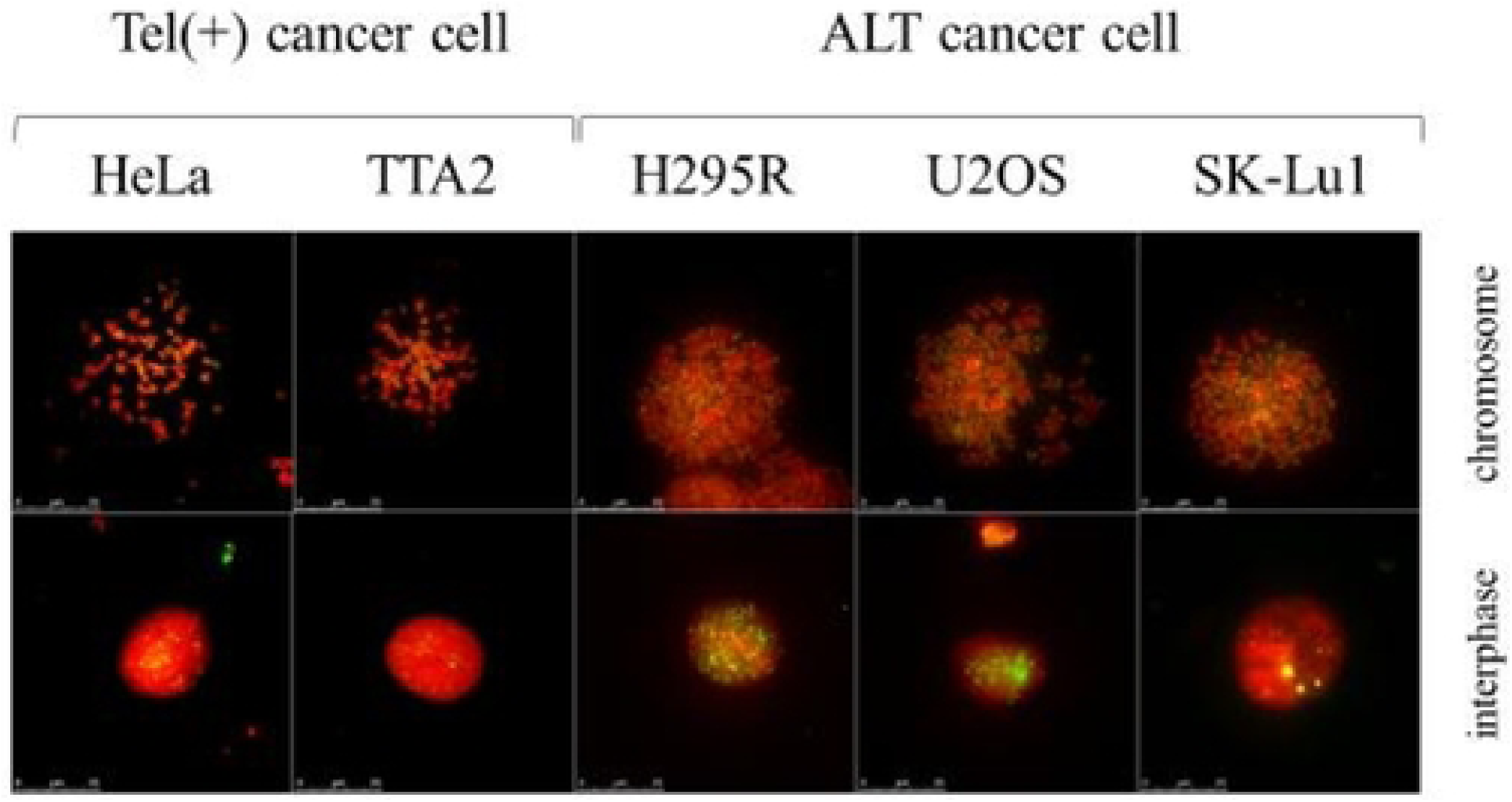
Telomeric FISH of chromosomes and interphase nuclei. Telomeric FISH of chromosomes (upper panel) and interphase nuclei (bottom panel). HeLa and TTA2 cells are Tel (+) cancer cells, and H295R, U2OS and SK-Lu1 cells are ALT cancer cells. Telomere signals were heterogeneous in interphase nuclei of ALT cancer cells, and giant telomeric signals were observed

### Distribution of telomeric FISH IVs in Tel (+) cancer cells and ALT cancer cells

Telomeric FISH IVs were calculated for 80 cells at each of the chromosomal ends and interphase nuclei to evaluate their distributions (Fig 2). Only ALT cancer cells showed different distributions of telomeric sizes between chromosomes and interphase nuclei. In HeLa and TTA-2 cells, telomeric FISH IVs at chromosomal ends exhibited similar distributions to those of interphase nuclei. In H295R, U2OS, and SK-LU1 cells, telomeric FISH IVs of interphase nuclei were larger than those of chromosomes. The differences were statistically significant by Welch’s *t*-test (P < 0.05) for the three ALT cancer cells.

**Fig 2.**
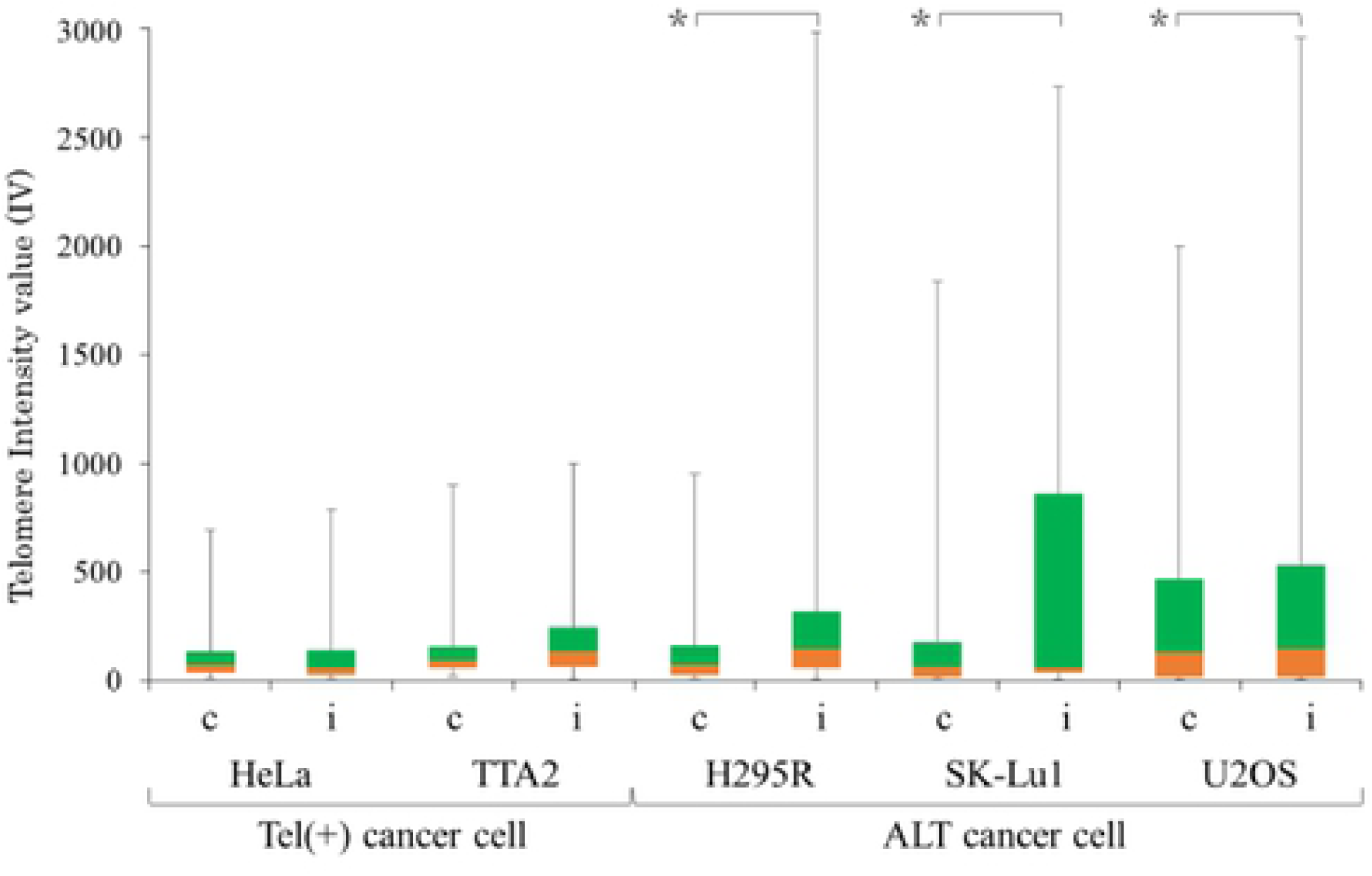
Distribution of telomeric IVs in Tel (+) cancer cells and ALT cancer cells. Distributions of telomeric FITC IVs in chromosomal specimens (c) and interphase nuclei (i) are shown in quartiles. Orange: first quartile median, Green: third quartile median. Distributions of telomeric IVs of interphase nuclei were significantly larger than those of chromosomes in ALT cancer cells (H295R, U2OS, SK-Lu1 cells) (*P < 0.05)

### Hierarchized telomeric FISH IVs of chromosomes and interphase nuclei

The telomeric FISH IVs of Tel (+) cancer cells and ALT cancer cells were stratified to compare chromosomes and interphase nuclei (Fig 3). In both chromosomal and interphase specimens of Tel (+) cancer cells, telomeric IVs were small and distributed in the range of 0–1000. In interphase nuclei of ALT cancer cells, larger telomeric signals were observed (greater than 1500) with a small peak at 2000–2500. In the chromosomal specimens, the majority of telomeric IVs were small and distributed in the range of 0– 1500.

**Fig 3.**
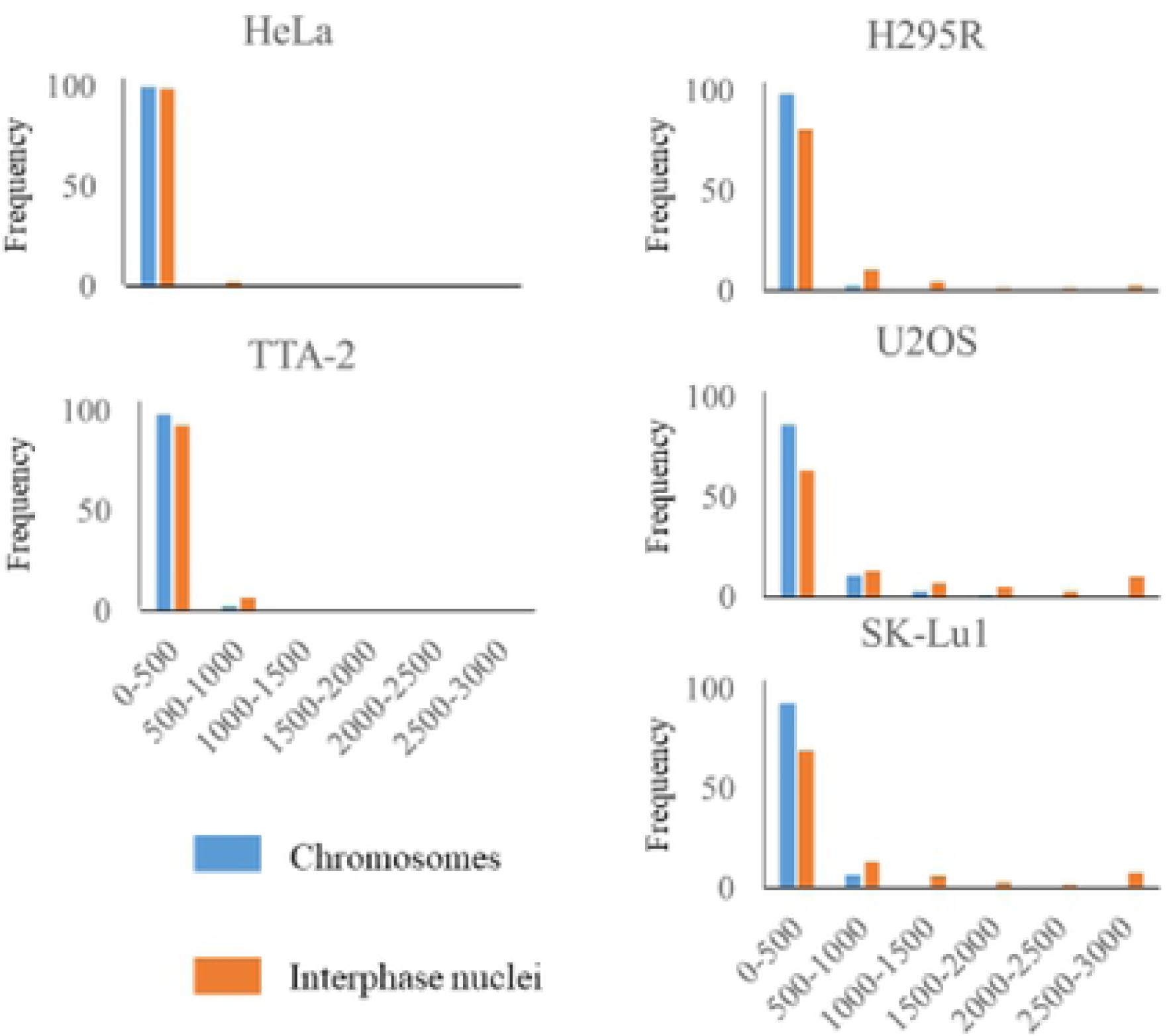
Hierarchized distributions of telomeric IVs of chromosomes and interphase nuclei. Telomeric IVs of chromosomes (blue bar) and of interphase nuclei (red bar) are demonstrated in a hierarchized graph for Tel (+) cancer cells and ALT cancer cells.

Giant telomeres that were greater than those at chromosomal ends were only observed in interphase nuclei of ALT cancer cells. GT-NBs specific to ALT cancer cells could be defined as larger than the maximum size of telomeres at chromosomal ends. Therefore, we could quantitatively define GT-NBs in each ALT cancer cell line. Consequently, GT-NBs of H295R and SK-Lu1 were defined as telomeric FISH signals of 1000 IVs or greater (8.7% and 10.6%, respectively) and those of U2OS cells were of 2000 IVs or greater (12.5%). In addition, GT-NBs were two to three times larger than the maximum size at the chromosomal ends in the three cell lines.

### Telomeres per cell

We evaluated telomeric signals per cell in chromosomes and interphase nuclei of each cell line. In H295R, U2OS, and SK-LU1 cells, the signal numbers were significantly lower in interphase nuclei than in chromosomes (P > 0.05), although there was no difference in HeLa or TTA2 cells (Fig 4) based on Welch’s *t*-tests. The second and third interquartile ranges for chromosomes and interphase nuclei did not overlap in H295R, U2OS, and SK-Lu1 cells, although they overlapped in HeLa and TTA-2 cells.

**Fig 4.**
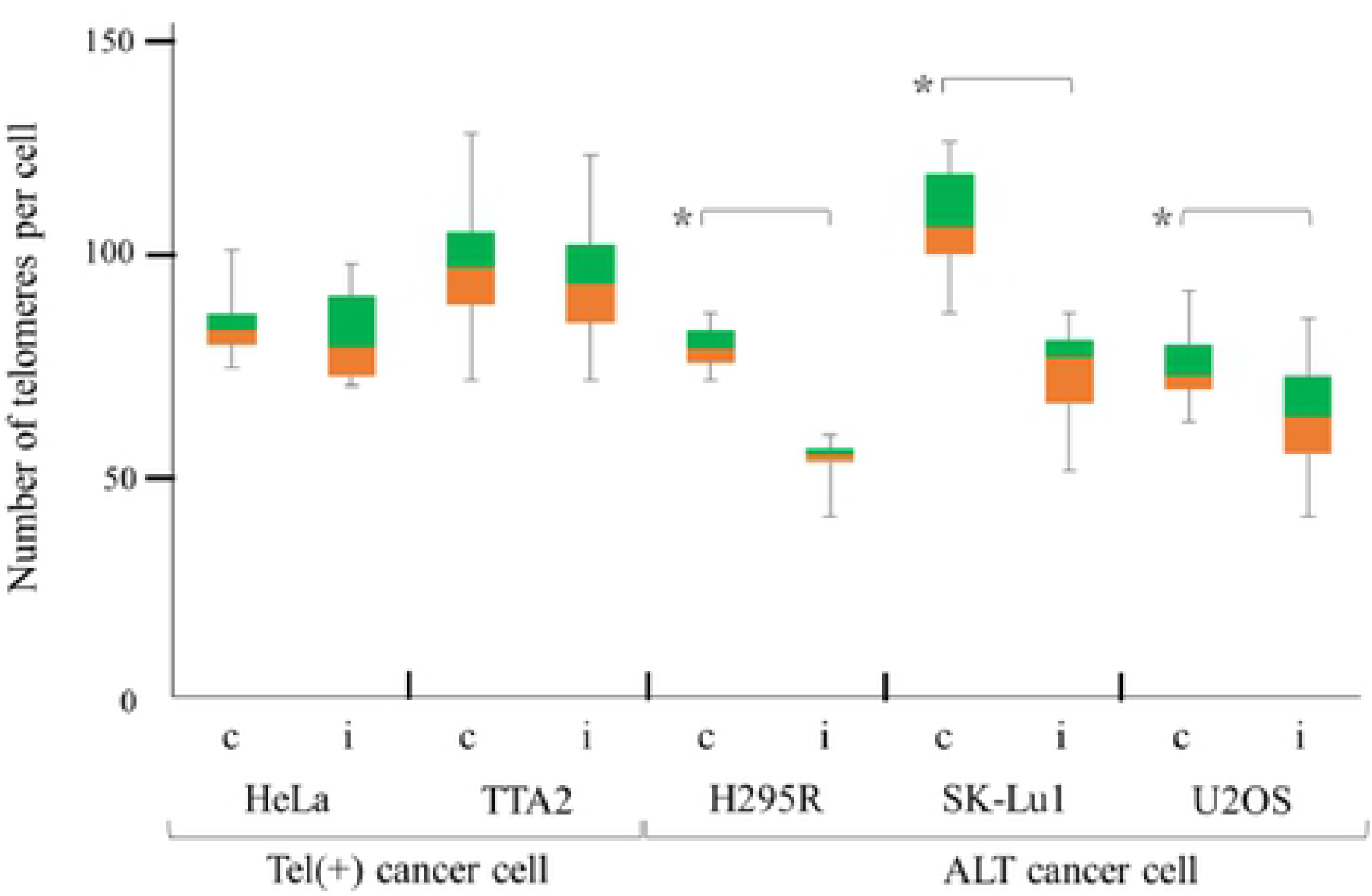
Telomeres per cell in chromosomes and interphase nuclei. Telomere counts per cell of chromosomal specimen (c) and interphase nuclei (i) are shown in quartiles. Orange: first quartile-median, Green: median-the third quartile. Numbers in ALT cancer cells were significantly lower in interphase nuclei than in chromosomes (P < 0.05).

### Co-localization of GT-NBs and PML-NBs

To analyze the ultra-structures of GT-NBs co-localized with PML-NBs, we double-stained interphase nuclei of H295R cells with telomeric FISH and anti-PML-NB antibodies and captured super-resolution (SR) microscopic images as well as fluorescence microscopic images (Fig 5). We examined 25 randomly selected interphase nuclei of H295R cells by fluorescence microscopy, and GT-NBs were co-localized with PML-NB in all 25 cells. Furthermore, we analyzed the microstructures of GT-NBs of five representative cells by SR microscopy. To confirm GT-NBs on SR microscopic images, the double fluorescence microscopic images of identical cells were overlaid (Fig 5), since SR microscopy only demonstrates the centers of gravity of fluorescent signals. The merged images revealed the microstructural relationship of GT-NBs and PML-NBs (Fig 5). Most GT-NBs include two PML-NBs on both sides of the GT-NB centers.

**Fig 5.**
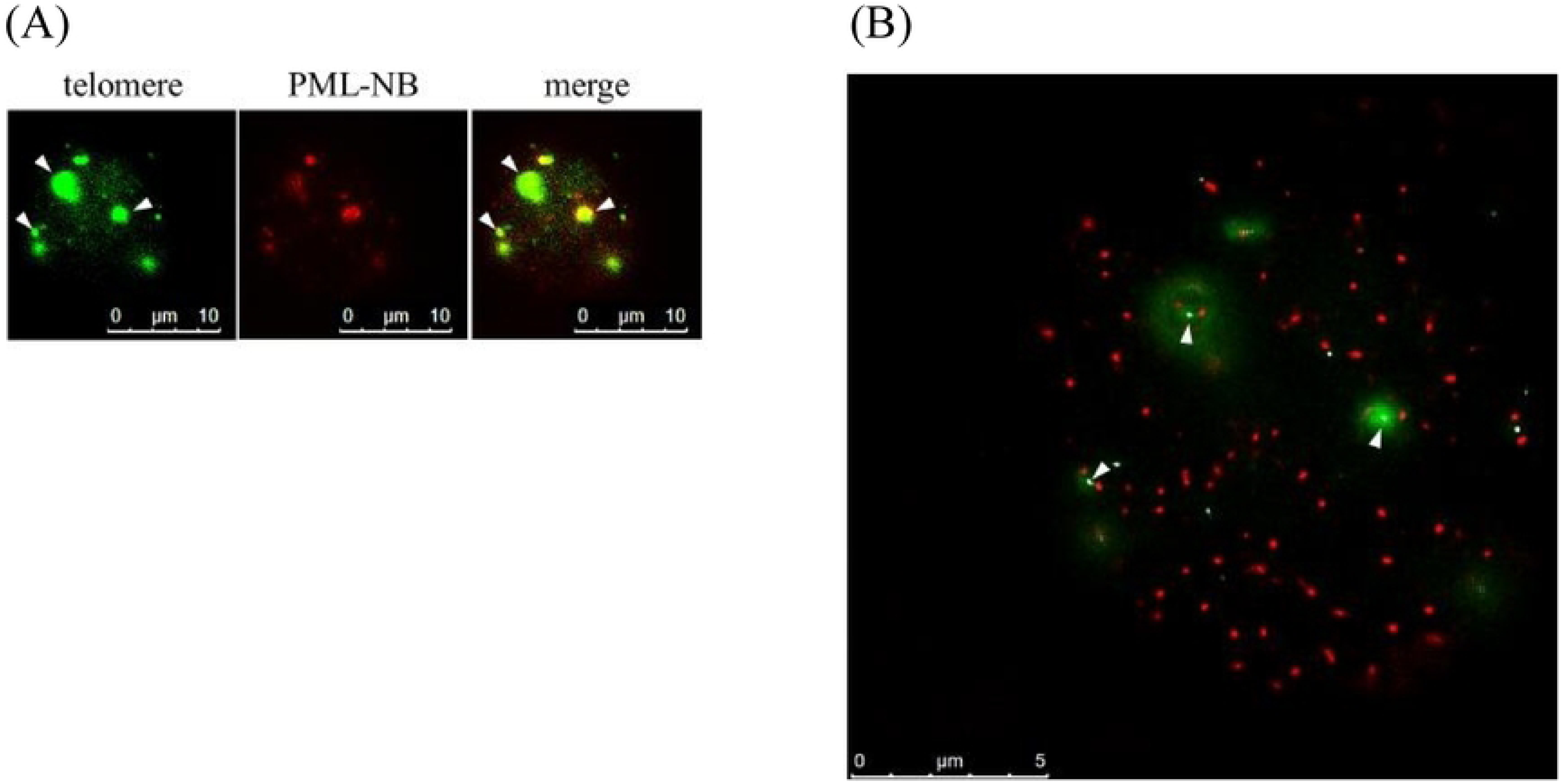
Fluorescence microscopic and super-resolution microscopic images of the co-localization of GT-NB and PML-NB. (A) Double-stained fluorescence microscopic images of an interphase nucleus of a H295R cell. left: telomeric FISH signals, middle: PML-NB signals, right: merged image. white arrowheads: GT-NBs (B) Super-resolution microscopic image combined with fluorescence microscopic image. GTNBs are shown as fuzzy green areas (fluorescence microscopic image), and they contain one central white spot (super-resolution microscopic image). Many PML-NBs are shown as red spots (super-resolution microscopy image). Within most GTNBs, two PML-NBs (red spots) are located on both sides of the center of the GT-NB (white spots)

## Discussion

The purpose of this study was to strictly define GT-NBs that are specifically observed in ALT cancer cells but not in Tel (+) cancer cells. It was possible to morphometrically define GT-NBs by their stronger signals in interphase nuclei than the maximum signals at metaphase chromosomal ends. In H295R ALT cancer cells, there were fewer telomeric signals at interphase nuclei than at chromosomes, but this difference was not detected in HeLa Tel (+) cancer cells. The reduction of telomeric signals can be explained by the fusion of telomeres in interphase nuclei of ALT cancer cells. In other words, multiple telomeric DNAs at chromosomal ends aggregate and form GT-NBs in ALT cancer cells.

All GT-NBs co-localized with PML-NBs in ALT cancer cells, i.e., GT-NBs were ALT-associated PML-NBs (APBs). Two hypotheses have been proposed to explain the morphogenesis of APBs: (1) PML-NBs enclose telomeric DNA by forming a shell and (2) PM-NBs are sites for the aggregation and interaction of telomeric DNAs. However, the resolution of fluorescence microscopy is too low to evaluate these hypotheses. In this study, we used super-resolution microscopy to analyze the ultra-structures of APBs and found that one or two PML-NBs existed within a GT-NB in ALT cancer cells. These results support the latter hypothesis. PML-NBs ensure physical proximity for interactions among telomeric DNAs. Furthermore, the observation of multiple PM-NBs within GT-NBs suggests that multiple interactions between telomeric DNAs occur within GT-NBs of ALT cancer cells.

Telomere elongation by the ALT mechanism is suggested by telomeric homologous recombination. This homologous recombination (HR) mechanism involves two process. First, the 3′ end of the protruding telomere sinks into the double-stranded portion of the other chromosome end. This process results in the formation of an HR intermediate structure called a Holliday junction. Next, the replication and extension of telomeric DNA occurs. Telomeres at chromosomal ends aggregate and the structure can be identified as a GT-NB by telomeric FISH. Therefore, our findings support the notion that GT-NBs of interphase of ALT cancer cells represent an intermediate structure in homologous recombination in the ALT mechanism. Though speculative, this is supported by the role of PML-NB in cells. Internal PML-NB presents various proteins and exhibits a variety of functions. Therefore, proteins function in DNA replication and transcription, apoptosis, the regulation of cell growth, and assistance with homologous recombination. In PML-NBs, double-stranded DNA repair occurs by homologous recombination after DNA damage. PML-NBs within GT-NBs facilitate intermediate structure formation by the homologous recombination of telomeric DNA.

In has recently been reported that ALT cancer cells possess telomeric sequences containing the intranuclear receptor-related TCAGGG sequence throughout their genomes. According to one such report, giant telomeric signals disappear by knocking down an intracellular receptor, NuRD-ZNF827, in ALT cancer cells. Therefore, ALT-specific GT-NBs suggested that the TCAGGG sequence is involved in nuclear receptor activity. Further analyses of GT-NB formation will elucidate the mechanism underlying ALT.

## Acknowledgments

We thank Ms. Ayumi Sumiishi, Ms. Kaoruko Kojima, and Ms. Namiko Kondo for technical support.

